# Ecological and evolutionary insights into pathogenic and non-pathogenic rice associated *Xanthomonas*

**DOI:** 10.1101/453373

**Authors:** Kanika Bansal, Samriti Midha, Sanjeet Kumar, Amandeep Kaur, Ramesh V. Sonti, Prabhu B. Patil

**Author notes:** Institute of Infection and Global Health, University of Liverpool, UK. Department of Archaeogenetics, Max Planck Institute for the Science of Human History, Jena, Germany (Sanjeet Kumar). Corresponding author, Address correspondence to Prabhu B. Patil.

## Abstract

*Xanthomonas oryzae* is a devastating pathogen of rice worldwide, however, *X. sontii* and *X. maliensis* are its non-pathogenic counterparts from the same host. So far, these non-pathogenic isolates were overlooked due to their less economic importance and lack of genomic information. We have carried out detailed ecological and evolutionary study focusing on diverse lifestyles of these strains. Phylogenomic analysis revealed two major lineages corresponding to *X. sontii* (ML-I) and *X. oryzae* (ML-II) species. Interestingly, one of the non-pathogenic *Xanthomonas* strains belonging to *X. maliensis* is intermediary to both the major lineages/species suggesting on-going diversification and selection. Accordingly, pangenome analysis revealed large number of lifestyle specific genes with atypical GC content indicating role of horizontal gene transfer in genome diversification. Our comprehensive comparative genomic investigation of major lineages has revealed that impact of recombination is more for *X. sontii* as compared to *X. oryzae*. Acquisition of type III secretion system and its effectome along with a type VI secretion system also seem to have played a major role in the pathogenic lineage. Other known key pathogenicity clusters or genes like biofilm forming cluster, cellobiohydrolase and non-fimbrial adhesin (*yapH)* are exclusive to pathogenic lineage. However, commonality of loci encoding exopolysacharide, *rpf* signalling molecule, iron-uptake, xanthomonadin pigment, etc. suggests their essentiality in host adaptation. Overall, this study reveals evolutionary history of pathogenic and non-pathogenic strains and will further open up a new avenue for better management of pathogenic strains for sustainable cultivation of a major staple food crop.

## Introduction

*Xanthomonas* is a complex group of bacteria that is primarily known for its pathogenic lifestyle causing infection in 125 monocots and 268 dicots (Leyns, De Cleene et al. 1984, Hayward 1993, Vauterin, Yang et al. 1996, Chan and Goodwin 1999) with remarkable host and tissue specificities. Among the 34 species of *Xanthomonas* (Parte 2018), *Xanthomonas oryzae* is solely known to be associated with rice pathogenicity which has further two pathovars on the basis of rice plant tissue infected as *X. oryzae* pv. oryzae and *X. oryzae* pv. oryzicola, infecting vascular and parenchyma tissues, respectively (NIÑO□LIU, Ronald et al. 2006). However, *X. sontii, X. maliensis* and some strains of *X. sacchari*, other *Xanthomonas sp.* were found to be associated with healthy rice seeds or leaves (Vauterin, Yang et al. 1996, Fang, Lin et al. 2015, Triplett, Verdier et al. 2015, Bansal, Kaur et al. 2019). Rice pathogenic *Xanthomonas* are widely studied, however, their non-pathogenic counterparts isolated from healthy plants are vastly overlooked. These non-pathogenic bacterial populations may be coevolving with the plant and their systematic study might give insights into the evolution of pathogenic counterparts. With the advent of the genomics era and renewed interest in bacterial ecology and evolution, non-pathogenic bacterial strains are now of deeper interest.

Extensive research is underway to identify the genetic features fundamentally required for pathogenicity and virulence potential of *Xanthomonas*. This work has aided in the identification of virulence genes and pathogenesis-related clusters in their genomes (Van Sluys, Monteiro-Vitorello et al. 2002, Toth, Pritchard et al. 2006). Protein secretion is a fundamental determining feature for pathogenic or symbiotic interactions as well as for inter-bacterial competition. There are six types of bacterial secretion systems, out of which five are present in pathogenic *Xanthomonas oryzae* strains (Ray, Rajeshwari et al. 2002, Büttner and Bonas 2010, Souza, Andrade et al. 2011, Pruitt, Schwessinger et al. 2015). *Xanthomonas* sp. undergo different stages in the infection process including colonizing leaf surfaces as epiphytes, invasion of the host, etc. Host plant has immune receptors for monitoring their extracellular and intracellular environment for the presence of bacteria. The *rax* gene cluster encodes a T1SS and a secreted sulfated peptide called RaxX which is recognized by rice receptor kinase XA21(Pruitt, Schwessinger et al. 2015). Further, ability to adhere to the host surface is important for successful infection. Among non-fimbrial adhesins, *xadA, xadB* help in attachment to the leaf surface and *yapH* aids in colonization of bacteria in xylem vessels (Das, Rangaraj et al. 2009). Amongst the fimbrial adhesins, the type IV pilus is also reported to help in attachment to the host (Juhas, Crook et al. 2008). *In planta* experiments indicate that *xadA, xadB* and *yapH* mutants show deficiency in the initial stages of infection like leaf attachment and entry (Das, Rangaraj et al. 2009).

*Xanthomonas* strains are known to have two representative T2SS gene clusters *xps* and *xcs*. Out of these, *X. oryzae* pv. oryzae and *X. oryzae* pv. oryzicola are equipped with one (*xps*) and *X. campestris* pv. campestris, *X. citri* pv. citri have both the systems (Szczesny, Jordan et al. 2010). *X. oryzae* uses the *xps* system for secreting hydrolytic enzymes to degrade rice cell wall, a process that appears to be crucial for pathogenesis (Jha et al., 2007). Further, the T3SS is one of the key pathogenicity factors in Gram-negative bacterial pathogens and is used to inject effectors into the host plant (Ghosh 2004). The T4SS is involved in transporting effectors and nucleoprotein complexes to the extracellular milieu or directly into the cytoplasm of other cells (Juhas, Crook et al. 2008). The T4SS is known to be involved in horizontal gene transfer, thus contributing to the genome plasticity and evolution of these bacteria. T4SS is absent among *X. oryzae* strains, though it is present in *X. albilineans, X. citri, X. campestris* and *Stenotrophomonas maltophilia* (Souza, Andrade et al. 2011).T6SS is a recently discovered secretion system described in human pathogens *Pseudomonas aeruginosa* and *Vibrio cholera* (Mougous, Cuff et al. 2006, Pukatzki, Ma et al. 2006). T6SS proteins are structurally and evolutionary related to phage proteins and contribute towards virulence, symbiosis, inter-bacterial interactions etc. (Sarris, Skandalis et al. 2010, Schwarz, West et al. 2010, Bernal, Llamas et al. 2018).

In response to ambient conditions, bacteria have evolved regulatory systems to regulate expression of pathogenicity related genes. For instance, *rpf* gene cluster involved in synthesis, detection and signal transduction of diffusible signal factor (DSF) is known to regulate pathogenicity in *Xanthomonas* (Barber, Tang et al. 1997, Wang, He et al. 2004, He, Zhang et al. 2007, Ryan, Fouhy et al. 2007, He and Zhang 2008). Further, for recognition and signal transduction, > 50 two component systems are predicted in *Xanthomonas* and, RavA/RavR, RpfC/RpfG, RaxH/RaxR, ColR/ColS and key response regulator HrpG are known to contribute to the virulence (Büttner and Bonas 2010). PhoPQ regulates expression of *hrpG*, which activates *hrpX* for induction of HrpG-induced genes (Tsuge, Nakayama et al. 2006, Lee, Jeong et al. 2008, Feng, Song et al. 2009) (Wengelnik and Bonas 1996, Tsuge, Terashima et al. 2005, Koebnik, Krüger et al. 2006).

Iron uptake is always a limiting factor in bacterial growth and affect virulence in case of animal and plant pathogenic bacteria (Cody and Gross 1987, Meyer, Neely et al. 1996, Bearden, Fetherston et al. 1997, Franza, Mahé et al. 2005, Lawlor, O’Connor et al. 2007). Iron regulated *xssABCDE* (*Xanthomonas* siderophore synthesis), *xsuA* (*Xanthomonas* siderophore utilization), *feo* (ferrous iron transporter) and *fur* (ferric uptake regulator) are involved in biosynthesis and regulatory roles of siderophores (Pandey and Sonti 2010) (Subramoni, Pandey et al. 2012). Genome-wide expression studies of *xibR* (*Xanthomonas* iron binding regulator) mutant have revealed that it regulates iron homeostasis in response to changing iron availability in environment (Pandey, Patnana et al. 2016). In addition to these, *Xanthomonas* species is known to produce extracellular polysaccharide (EPS) via *gum* gene cluster, whose mutants have been largely studied for effects on virulence (Chou, Chou et al. 1997, Katzen, Ferreiro et al. 1998, Dharmapuri and Sonti 1999). Also, xanthomonadin, a characteristic feature of *Xanthomonas* which is encoded by the *pig* cluster provides protection against photo oxidative damage (Stephens and Starr 1963, Goel, Rajagopal et al. 2002) (Goel, Rajagopal et al. 2001, Cao, Wang et al. 2018).

A comprehensive genomic investigation of pathogenic *Xanthomonas* with non-pathogenic *Xanthomonas* strains from the same host can help in identifying the functions acquired, lost or modified during adaptation to a particular lifestyle by a bacterial strain. As the genome of a successful pathogen is shaped by genetic, ecological and evolutionary factors, this comparative study may give a clearer picture of the evolution of bacterial pathogenesis on rice in particular and plants in general. Such a study can also help in developing newer methods of disease control and putative genomic markers for proper diagnosis/identification.

## Results

### Strains used in the study

For the present study, strains known to be non-pathogenic towards rice i.e. African strain LG27592; Chinese strains LMG12459, LMG12460, LMG12461 and American strain LMG12462 (Vauterin, Yang et al. 1996, Triplett, Verdier et al. 2015) were procured from the Belgian Co-ordinated Collections of Micro-organisms/ Laboratory of Microbiology Gent Bacteria Collection (BCCM/LMG). Further, genome sequences of seven strains available on NCBI GenBank database were retrieved (three *X. sontii* strains PPL1, PPL2, PPL3, American strains SHU166, SHU199, SHU308 and Chinese strain *X. sacchari* R1). These strains are isolated from surface-sterilized rice seeds. *X. sontii* strains are reported to be non-pathogenic towards rice, however, pathogenicity status of remaining isolates is unknown (Fang, Lin et al. 2015, Bansal, Kaur et al. 2019).

Further, based on a previous study from our group, there are five distinct lineages of Asian *X. oryzae* pv. oryzae strains (Midha, Bansal et al. 2017). We have selected one representative strain from each of the lineages (BXO1, IXO222, XOOP, XOOK and IXO599 from L-I, L-II, L-III, L-IV and L-V respectively) as the pathogenic counterpart. We also included representative strains of different geographic locations (BXO1 from India; AXO1947, CIX44 from Africa; PXO86 from the Philippines; X8-1A from US; YM15 from China and CFBP2286 from Malaysia). In addition, we have also included one *X. oryzae* pv. oryzicola strain (BLS256).

### In-house whole genome sequencing and assembly

Whole genome sequencing of five non-pathogenic isolates was carried out on an in-house Illumina MiSeq platform. The high quality de novo assembly of the Illumina reads resulted in genomes with coverage ranging from 155x to 202x and with N50 values of 50.7 kb to 77.5 kb (table 1). The genome size for all the NPX strains was found to be approximately 5 Mb, which is similar to the pathogenic *Xanthomonas* genus, suggesting no large-scale reductive evolution in these non-pathogenic strains.

**Table 1.** Genome statistics of genomes sequenced in the present study.

| S. N. | Strain Name | Completeness/<br>Contamination | Fold (X) | N50 (Kb) | Contigs | Genome Size | GC(%) | CDS | rRNA +tRNA | Isolation source | Accession No. |
| --- | --- | --- | --- | --- | --- | --- | --- | --- | --- | --- | --- |
| 1 | *X. maliensis* LMG27592 | 96.74/0.30 | 155 | 77.5 | 141 | 5.2 | 66.1 | 4369 | 3+53 | Rice leaves | NSET |
| 2 | *Xanthomonas* sp. LMG12459 | 98.17/0.86 | 184 | 59.3 | 143 | 4.8 | 68.8 | 4055 | 3+51 | Rice seeds | NSGT |
| 3 | *Xanthomonas* sp. LMG12460 | 96.58/0.00 | 180 | 50.7 | 178 | 4.7 | 69 | 3934 | 4+51 | Rice seeds | NKUH |
| 4 | *Xanthomonas* sp. LMG12461 | 98.76/0.64 | 202 | 71.8 | 134 | 4.8 | 68.9 | 4060 | 3+51 | Rice seeds | NQYT |
| 5 | *Xanthomonas* sp. LMG12462 | 97.64/0.52 | 176 | 76 | 136 | 4.8 | 68.9 | 4062 | 3+52 | Rice seeds | NMPP |

### Phylogenomic analysis of the non-pathogenic and pathogenic isolates

Phylogenomics clearly depicted that all the rice associated non-pathogenic strains (except *X. maliensis*) clubbed with the *X. sontii* forming the major lineage ML-I (figure 1). Eventhough, *X. sacchari* R1 was classified as *X. sacchari*, but in our analysis, it clubbed with ML-I i.e. *X. sontii* and not with *X. sacchari* CFBP4641 (T). All pathogenic strains (*X. oryzae*) formed another major lineage ML-II. Peculiarly, *X. maliensis* being a non-pathogenic strain, was phylogenomically close to pathogenic lineage. For phylogenomic analysis, *Stenotrophomonas maltophilia* ATCC13637 (T) was used as an outgroup. Further, since one of the strains among the NPX (*X. sacchari* R1) was classified in *X. sacchari*, type strains of *X. sacchari* and its closest relative *X. albilineans* were also included in the analysis (López, Lopez-Soriano et al. 2018).

**Figure 1:**
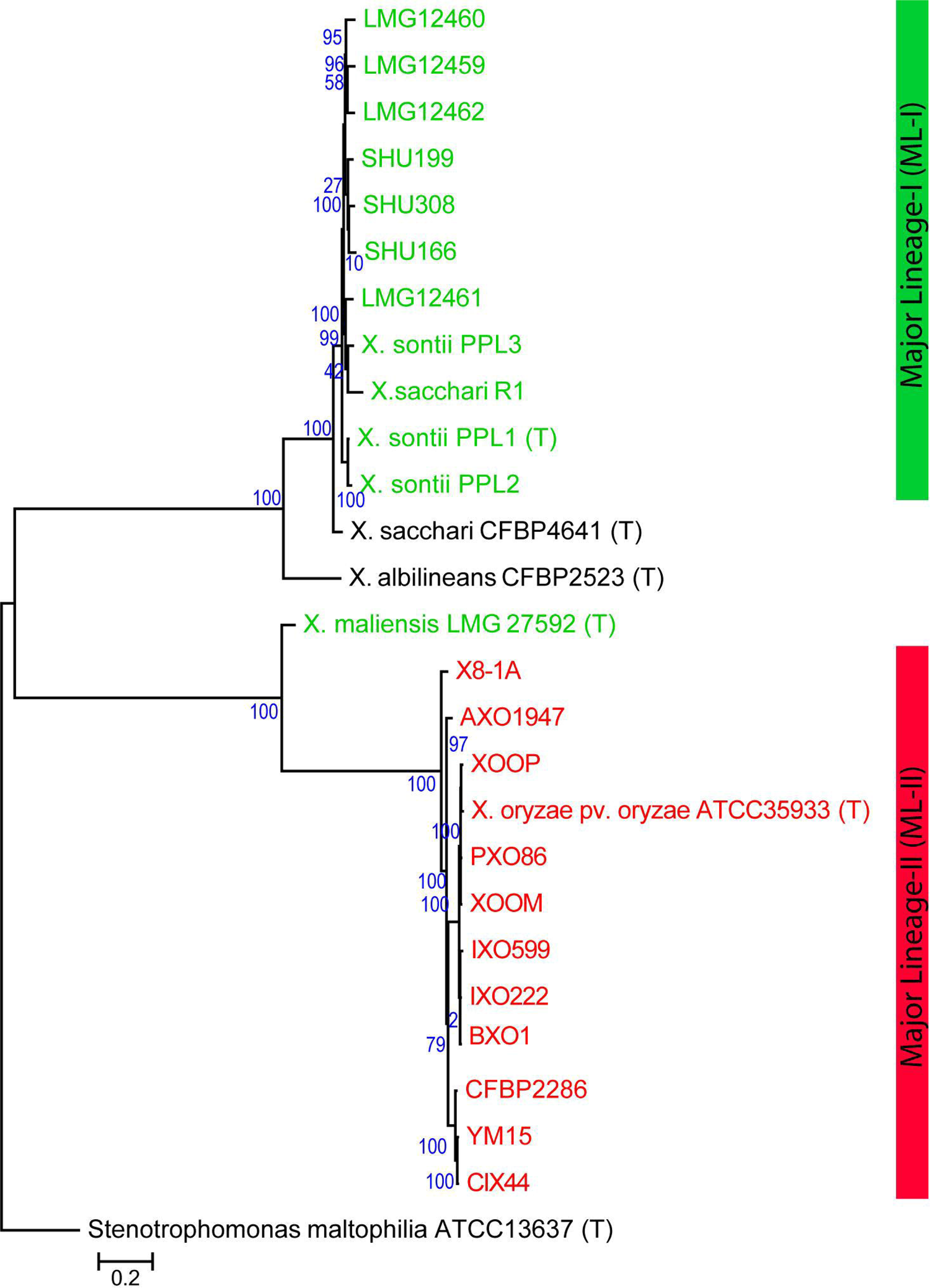
PhyloPhlAn tree of all rice associated strains. Rice non-pathogenic isolates are in green and pathogenic in red color. Here, strains of ML-I and ML-II are designated. *Stenotrophomonas maltophilia* ATCC13637 (T) was used as outgroup and other type strains are designated as (T).

### Taxonogenomic status of rice seed associated strains

To explicitly assess taxonomic status of ML-I and *X. maliensis* strains, we used orthoANI values, to define species status (figure 2). The type strains of *X. sontii, X. sacchari* and *X. albilineans* were also included in this analysis. OrthoANI uses species delineation cut off of 96% similarity (Richter and Rosselló-Móra 2009). All ML-I isolates were having ANI values >96.2% with *X. sontii* and that of around 93% with *X. sacchari* CFBP4641 (T) and around 83% with *X. albilineans* CFBP2523 (T). Hence, they belong to *X. sontii*, here also, even *X. sacchari* R1 was found to belong to *X. sontii*, depicting its misclassification. Further, the variant *X. maliensis*is strain is already reported as a distinct species (Triplett, Verdier et al. 2015). Thus it can be concluded that, all of the rice associated strains were not belonging to *X. oryzae*. They belong to three different species i.e. *X. sontii* (ML-I), *X. oryzae* (ML-II) and *X. maliensis.*

**Figure 2:**
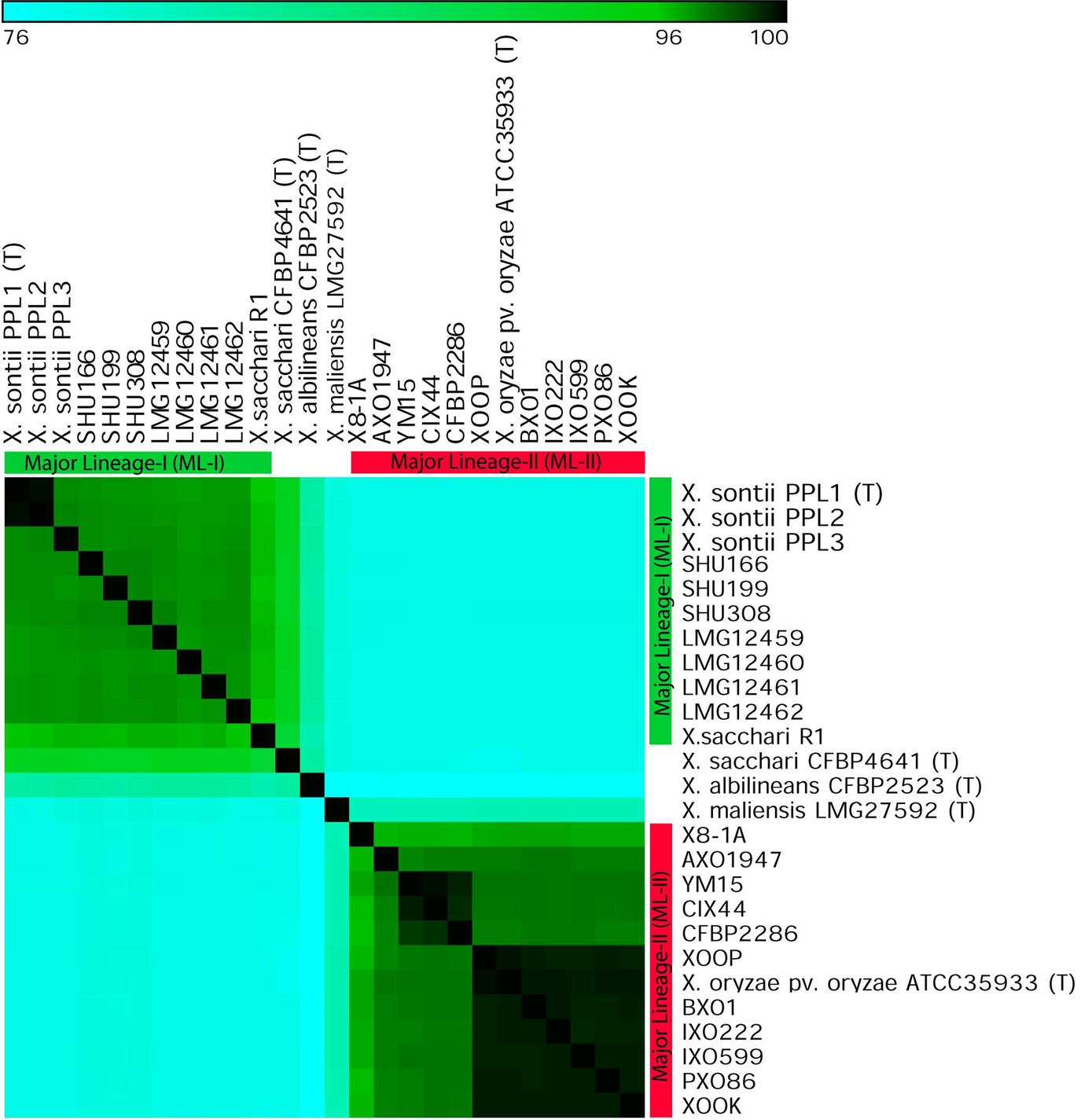
**OrthoANI of rice associated strains** including type strains of *Xanthomonas sacchari* CFBP4641 (T) and *Xanthomonas albilineans* CFBP2523 (T).

Since, *X. maliensis* was found to be more related to pathogenic isolates and not to non-pathogenic, this will be excluded from downstream analysis to remove the biasness. Hence, only major lineage strains (ML-I corresponding to *X. sontii* and ML-II corresponding to *X. oryzae*) are considered for further analysis.

### Recombination and mutation impact on lifestyle of bacteria

Recombination and mutations are major driving forces of bacterial evolution. To measure their impact on the major lineages, we carried out ClonalFrameML analysis. Interestingly, the output suggests that recombinational events are not very frequent amongst these clades of *Xanthomonas* strains. There is occurrence of around 5 and 12 mutational events for each recombinational event in *X. sontii* and *X. oryzae* respectively. However, even though there is low occurrence of recombination, it has more impact (r/m = 1.233) towards the evolution of strains in *X. sontii*. But, this scenario is not seen in the pathogenic strains, where the impact of recombination is only half that of mutation (r/m = 0.54).

### Distinct gene content among non-pathogenic and pathogenic isolates

To evaluate the gene-content wise relatedness of strains under study, we focused on their core and pan-genome content (figure 3). Here, core was least (1039) and pan was maximum (17221) for all strains (including all *Xanthomonas* strains under study and *Stenotrophomonas maltophilia*). Whereas, for all *Xanthomonas* strains (excluding *S. maltophilia*) under consideration, core increased (1291) and pan decreased (14231). Core and pan values were (1314, 13385) for *X. sontii* and *X. oryzae* strains taken together. Here, core values increased and pan values decreased dramatically, when *X. sontii* and *X. oryzae* strains were considered separately (2224, 6766; 2686, 9415). Interestingly, when *X. maliensis* was included in *X. sontii*, the core decreased and pan increased (1626, 8357), whereas inclusion of *X. maliensis* did not have such an effect on *X. oryzae* (2391, 10601). Pangenome analysis also emphasized that although a non-pathogenic isolate, *X. maliensis* is more related to *X. oryzae.*

**Figure 3:**
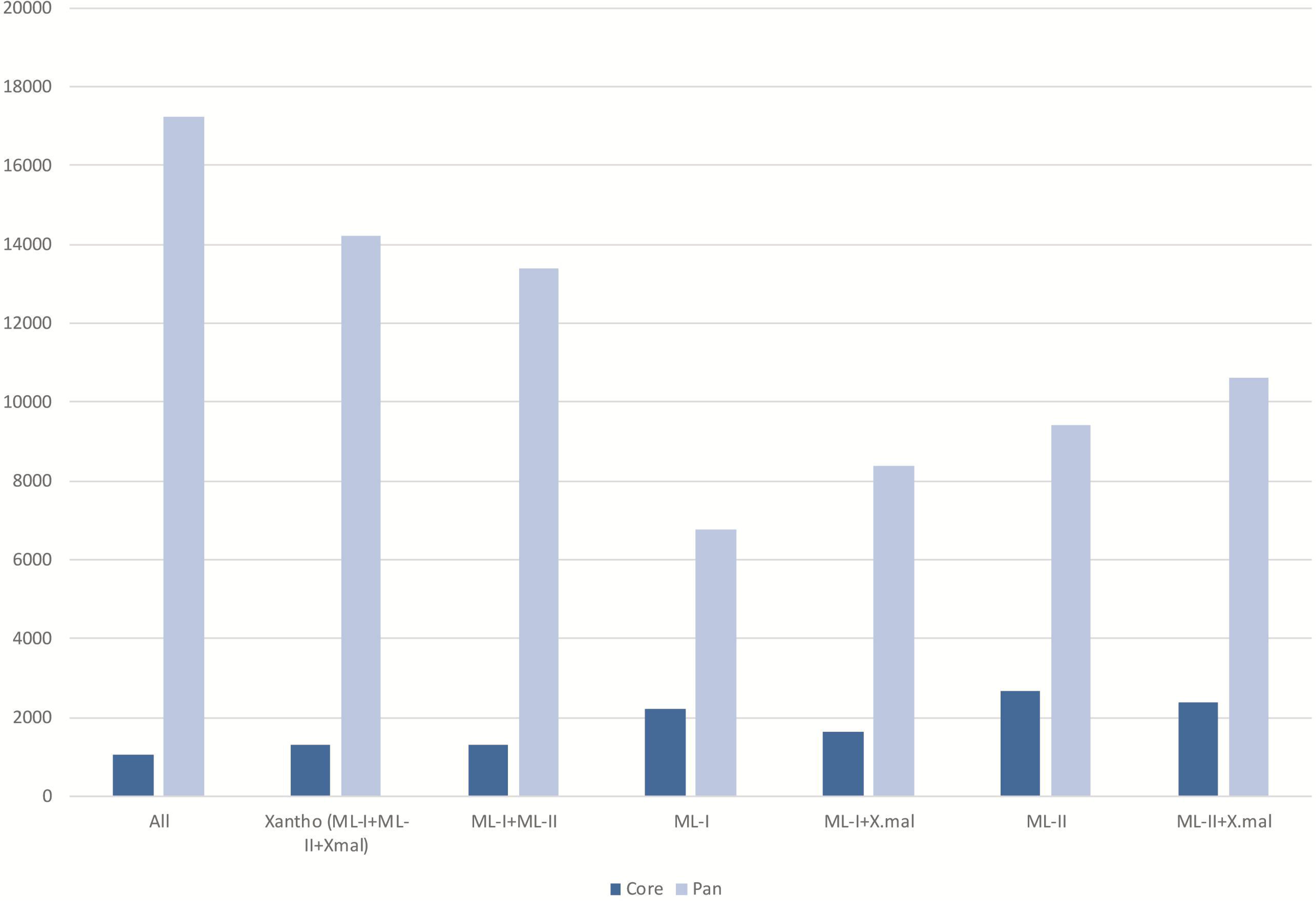
Core and pan genome analysis. Here, ML-I: *X. sontii*, ML-II: *X. oryzae* and Xmal: *X. maliensis.* Strains used for the analysis is indicated along x-axis and number of genes in y-axis.

Whether differences in lifestyles of the bacterial isolates are also reflected in the gene content? To address this, unique genes among different lifestyle bacteria were inspected manually. Unique genes to *X. sontii* were 686 (unique ML-I) and to *X. oryzae* were 583 (unique ML-II). Interestingly, *X. sontii* unique genes were having average GC content of 86% (with all genes from >67% GC) and *X. oryzae* unique genes GC content was 42% (with genes from both <61% and >67% GC). Whereas, typical GC content for *Xanthomonas* is within 64.5% ±2.5% range. Further, functional analysis of these genes was carried out. Among 25 COG classes, our data was represented in 17 classes while approx. 50% were having unknown function or hypothetical proteins (figure 4). Overall, genes related to metabolism (such as carbohydrate, lipid, inorganic ion and secondary metabolites metabolism biosynthesis and transport) and information storage and processing such as transcription were more in non-pathogenic as compared to pathogenic. While, genes unique to the pathogenic pool were more of cell wall/membrane/envelope biogenesis, motility related or intracellular trafficking, secretion and vesicular transport related genes. Whereas, genes related to amino acid and nucleotide transport and metabolism; replication, recombination and repair etc. were comparable among both the pathogenic and non-pathogenic strains.

**Figure 4:**
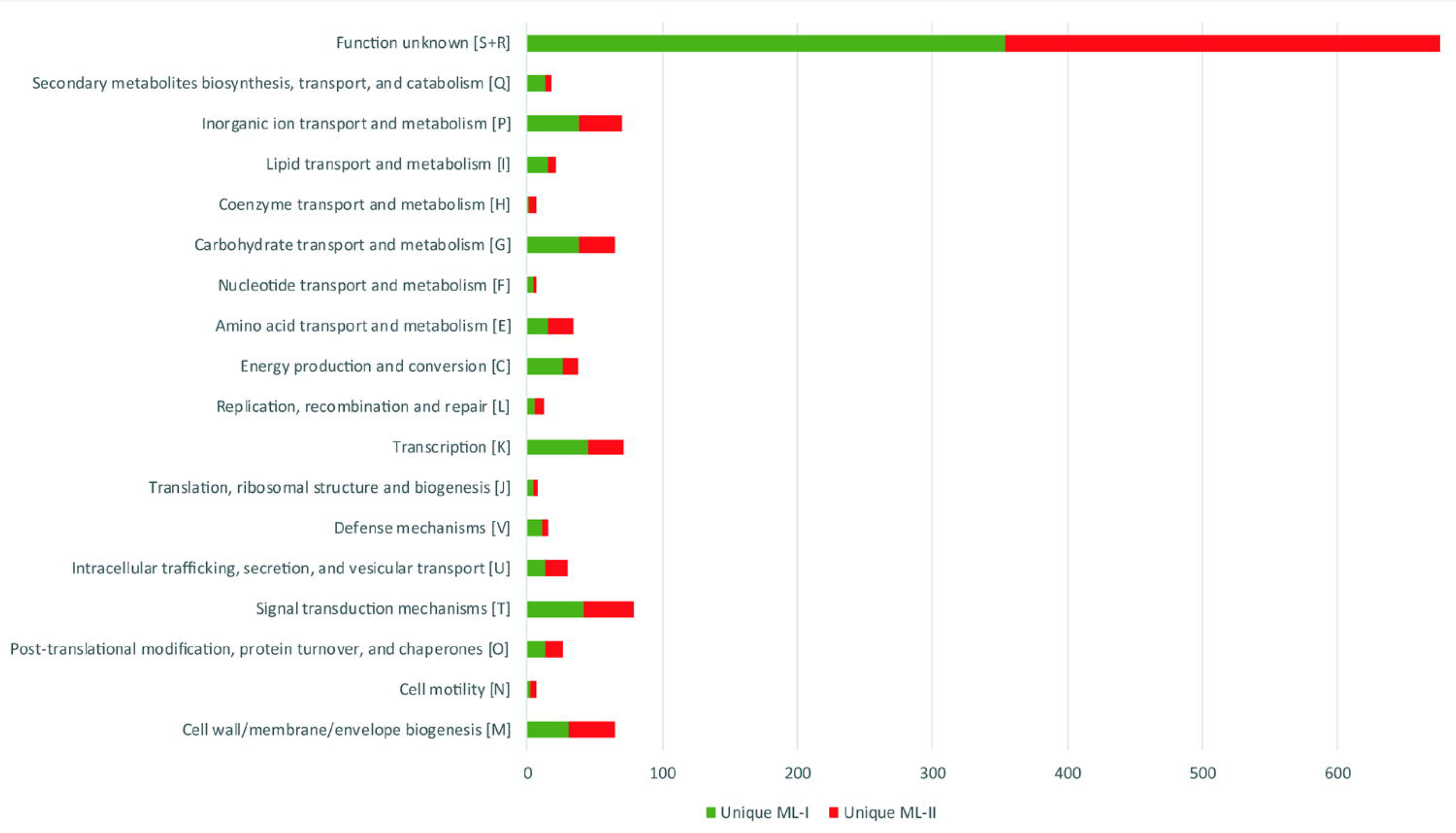
COG functional classification of unique genes. Unique genes to *X. sontii* (unique ML-I) are shown in green color and unique genes to *X. oryzae* (unique ML-II) are shown in red color.

Further inspection into the pangenome results were carried out by looking for the annotation of the genes unique to different sets of strains described above. Interestingly, unique *X. oryzae* were genes related to various types of secretion systems, PhoPQ-activated pathogenicity-related protein, etc. Strikingly, pan-genome analysis depicted that biofilm biosynthesis cluster (*pgaABCD*) is absent from *X. sontii* but is present in *X. oryzae* strains. Further, this was also confirmed experimentally by biofilm analysis (figure 5), *X. oryzae* BXO1 and *X. oryzae* pv. oryzicola displayed higher biofilm forming abilities than *X. sontii* strains. However, *X. maliensis* was found to display biofilm forming capability, which was later confirmed by genomic analysis, reaffirming its relatedness to *X. oryzae* and not to *X. sontii*.

**Figure 5:**
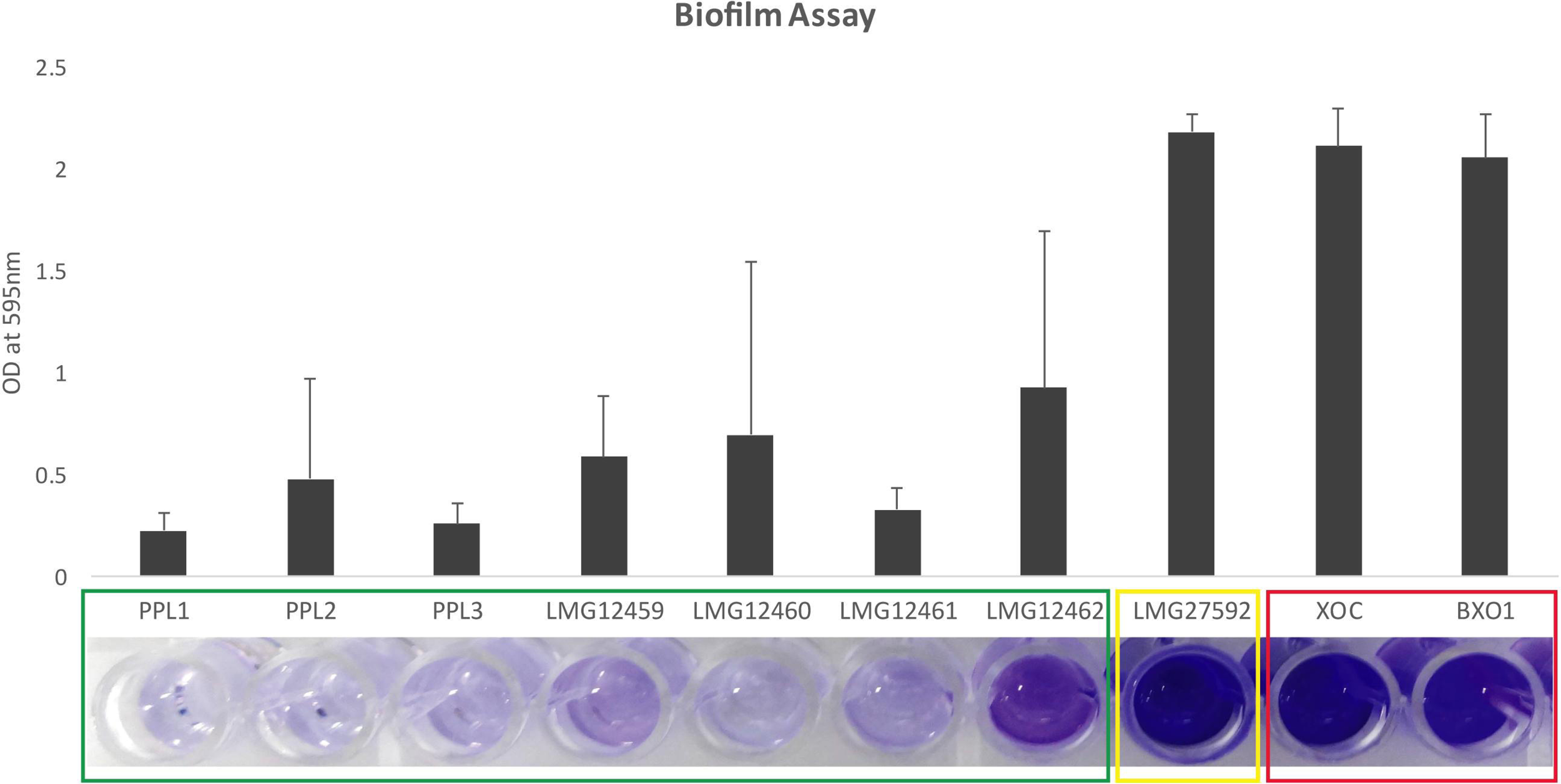
Biofilm assay: Ability of strains to form biofilms in borosilicate glass tubes determined using crystal violet staining. Here, *X. sontii, X. maliensis* and *X. oryzae* are represented in green, yellow and red boxes.

### Pathogenicity gene(s) and gene cluster(s) in rice associated *Xanthomonas*

*Xanthomonas* is a model phytopathogen and numerous genetic studies have identified major pathogenicity related gene(s) and gene cluster(s) (Büttner and Bonas 2010). We proceeded to examine the status of various well known *Xanthomonas* pathogenicity genes and gene clusters (supplementary table 1) amongst the three *Xanthomonas* species associated with rice. First, we focused on all the protein secretion systems, effectors produced by them and their regulators which could be playing a critical role in their interaction with the host (figure 6 (a), (b)). Interestingly, T1SS *raxX, raxST, raxA* showed variable or complete absence in non-pathogenic isolates, however, they were present in the pathogenic isolates. A single T2SS i.e. *xps*, is present in pathogenic and non-pathogenic strains, whereas *X. maliensis* has both of the secretion systems (*xps* and *xcs*).Type II system secreted effectors or cell wall degrading enzymes (CWDEs) which are well studied for *Xanthomonas* were examined in this study included cellulases (*cbsA, clsA, cel8A, cel9A*); xylanases *(xynA, xyn10A, xyn5A, xyn51A,xynB, xyn10B, xyn10C*); pectate lyases (*pel10A, pel1A, pel1B, pel1C, pel3A, pel4A, pel9A*); lipase (*lipA*) and *aguA* and *pglA* (Dow, Scofield et al. 1987, Ray, Rajeshwari et al. 2000, Rajeshwari, Jha et al. 2005, Potnis, Krasileva et al. 2011). Among the cellulases, *cbsA* is present only in pathogenic (except for *X. oryzae* pv. oryzicola CFBP2286, where it has a frameshift mutation) strains and was completely lacking in all the NPX isolates. Interestingly, *cel8A* and *cel9A* were present in all rice non-pathogenic but were absent in pathogenic strains (except for absence of *cel8A* in *X. maliensis*) and *clsA* is present in all isolates. However, xylanases and pectate lyases showed variations in their repertoire among pathogenic and non-pathogenic isolates. Among xylanases *xynA, xyn10A, xynB* and *xyn10B* are present in *X. oryzae* and *X. maliensis*; while *xyn10C* is present only in *X. maliensis*. The xylanases *xyn5A* and *xyn51A* are present in all the isolates (except for absence of *xyn5A* in CFBP2286, CIX44 and YM15). Further, *aguA* and *lipA* genes are present in all the isolates and *pglA* is present in *X. oryzae* and *X. maliensis*. In case of pectate lyases, *pel9A* is present in pathogenic and *X. maliensis* except for AXO1947, X8-1A and *pel1B* was present in all NPX strains except for *X. sacchari* R1. While, *pel1A* and *pel4A* are present in all the strains (except for absence of *pel1A* in X8-1A) and *pel3A, pel10A* and *pel1C* are not present in any of the strains under study. T3SS and its effectors are present in pathogenic strains and absent from all non-pathogenic strains (*X. sontii* and *X. maliensis*). While T4SS is only present in *X. sontii* isolates. Among adhesins (type V effectors), *yapH* is exclusive to pathogenic isolates, however, *xadA, xadB* and *pilQ* are present among all the isolates.

**Figure 6:**
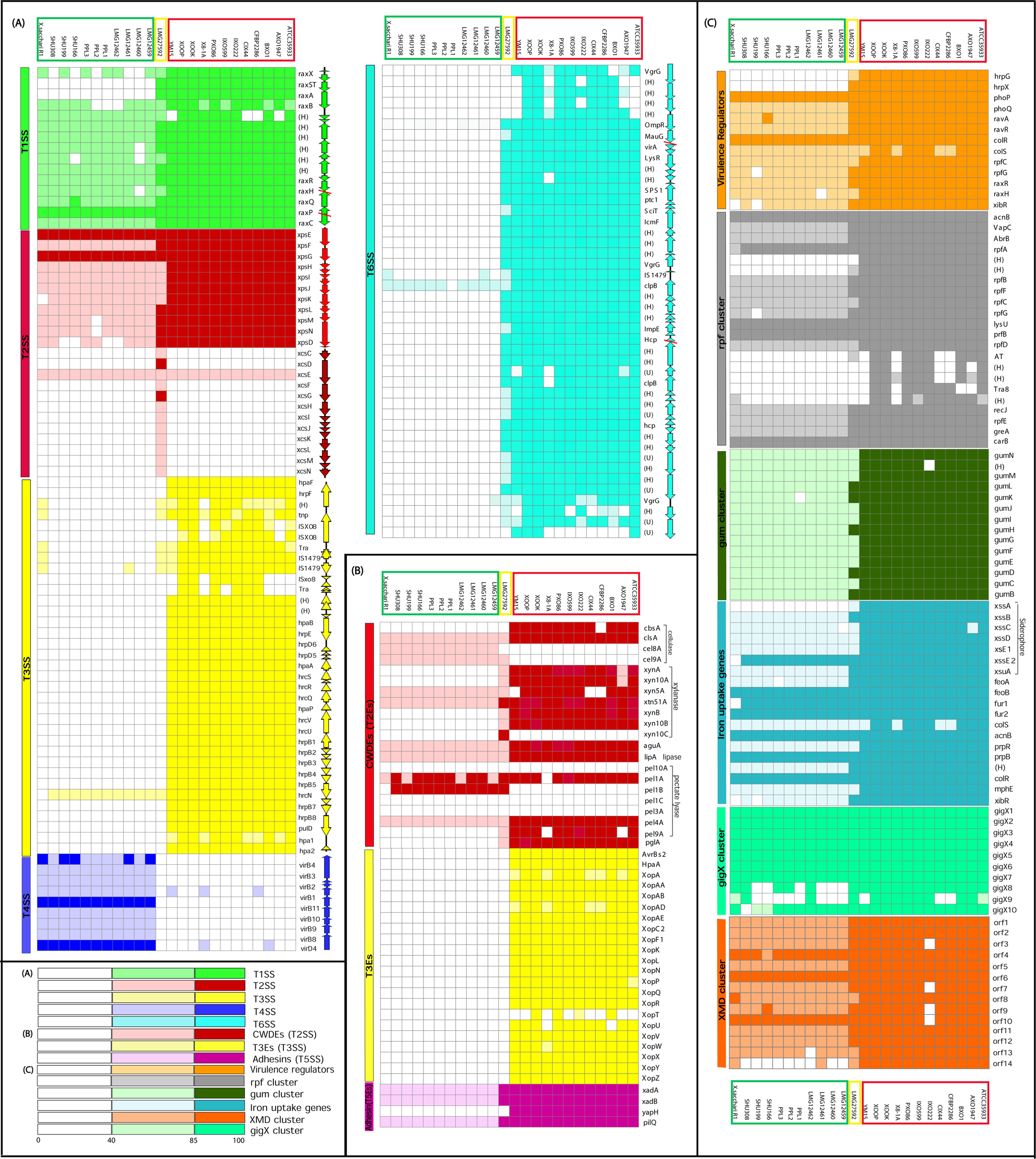
**Distribution of pathogenicity related gene(s) and gene cluster(s)** among *X. oryzae* (in red box), *X. sontii* (in green box) and *X. maliensis* (in yellow box). Different colors are corresponding to different clusters and the intensity of the color indicates level of identity with the query as depicted by the scale.

We also scanned for the presence of genes encoding virulence regulators (figure 6 (c)). The *rpf* cluster, is present in both groups of bacteria, although non-pathogenic isolates were having a diversified *rpf* cluster as compared to pathogenic isolates. Similarly, we looked for other regulators of virulence, most of them were present in all the isolates under study. However, *hrpG, hrpX* global regulators are present in *X. oryzae* and *X. maliensis* and absent in *X. sontii* strains. Other two component systems viz., RavA/RavR, RpfC/RpfG, RaxH/RaxR and ColR/ColS are present in all the isolates. Iron uptake (ferric and ferrous) in response to changing iron availability in environment is mediated via various two component systems, siderophores and *xibR*. Here, all strains showed presence of genes responsible for this, with diversified genes in *X. sontii* lineage, except for *xssA*, which is present only in *X. oryzae* and *X. maliensis*. Further, genes responsible for EPS production (*gum* cluster), yellow pigment production (xanthomonadin pigment) and flagellar glycosylation (*gigX* cluster) are present in all the isolates.

## Discussion

### Distinct evolutionary trajectories in rice associated *Xanthomonas*

Genome based studies are allowing researchers to obtain detailed and comprehensive insights into bacterial evolution. It can be clearly inferred from the present whole genome based analysis that large number of rice associated *Xanthomonas* strains formed two distinct major lineages (*X. sontii*, ML-I and *X. oryzae*, ML-II) associated with diverse lifestyles suggesting parallel evolution. This analysis also indicates that the non-pathogenic strains are not random associations with rice and they represent genuine adaptations of *Xanthomonas* to a non-pathogenic lifestyle on this host. In addition to these two rice associated species, *X. maliensis* is more related to pathogenic lineage (ML-II) eventhough its reported to have non-pathogenic lifestyle. This points out that this species has emerged independently to that of the *X. sontii* in close association with rice plant. The phylogenomic tree suggests that *X. sontii* and *X. oryzae* diverged a long time ago and also suggests a distinct rate and pattern of divergence taking place in these two species. Genome based studies not only allow us to understand bacterial diversity at the sequence level but also makes it possible to compare the gene content of pathogenic and non-pathogenic isolates. This can provide insights into evolution of different lifestyles of the bacteria from a common ancestor. Pangenome for both the lineages were open with large numbers of unique genes with unknown functions, highlighting the need for more functional genomics and genetic studies for a detailed understanding. However, atypical GC content of unique gene to the lineages were only in higher range (>67%) for *X. sontii* whereas, in both high and low ranges (<62% and >67%) for the pathogenic ones. This clearly depicts diverse sources and independent acquisition of unique genes and hence, supports distinct evolutionary trajectories led by pathogenic and non-pathogenic isolates. Moreover, pan genome size for pathogenic (9415) was relatively higher as compared to non-pathogenic (6766), which indicates pathogenic genomes to be dynamic as expected. Further, pangenome analysis allowed us to identify gene cluster for biofilm biosynthesis (*pgaABCD*) in all the strains of the *X. oryzae* and absent from *X. sontii*, suggesting importance of its function on evolution and emergence in pathogenic lineage of *Xanthomonas* in rice. Hence comparative genomic strategies have potential to identify novel genomic loci and functions associated with pathogenic life-style and also the potential to target them in management of pathogenic lineage.

*Xanthomonas* is a model bacterial pathogen to understand molecular aspects of host-pathogen interactions, virulence and pathogenicity (Lu, Patil et al. 2008, Büttner and Bonas 2010, Jacques, Arlat et al. 2016). Apart from pan-genome analysis, the present study has provided novel insights into the status and evolution of well characterized pathogenicity associated genes/gene clusters in two evolutionarily distinct and ecologically related isolates having diverse host-microbe interactions (pathogenic and non-pathogenic) (Price, Dehal et al.) (figure 7). For example, the hallmark of pathogenicity of *Xanthomonas* is T3SS and its effectors (White, Potnis et al. 2009) which pathogenic strains seem to have acquired by the ancestor of the pathogenic lineage (ML-II) after the divergence of pathogenic and non-pathogenic lineages and have played a primary role in their emergence and pathogenicity. T6SS, HrpG, HrpX regulons known to be critical for pathogenicity of *Xanthomonas* are present in all *X. oryzae* strains and even in *X. maliensis* as well, but are not present in *X. sontii*. It is consistent with previous reports of their importance for pathogenicity and it was acquired by common ancestor of *X. oryzae* and *X. maliensis*, i.e. after diversion of *X. sontii* during the evolution. Conversely, presence of T4SS in *X. sontii* suggests is importance in emergence and success of major lineage of non-pathogenic strains. In this case, it seems to be acquired by ancestor of *X. sontii* after divergence of *X. oryzae* and *X. maliensis*.

**Figure 7:**
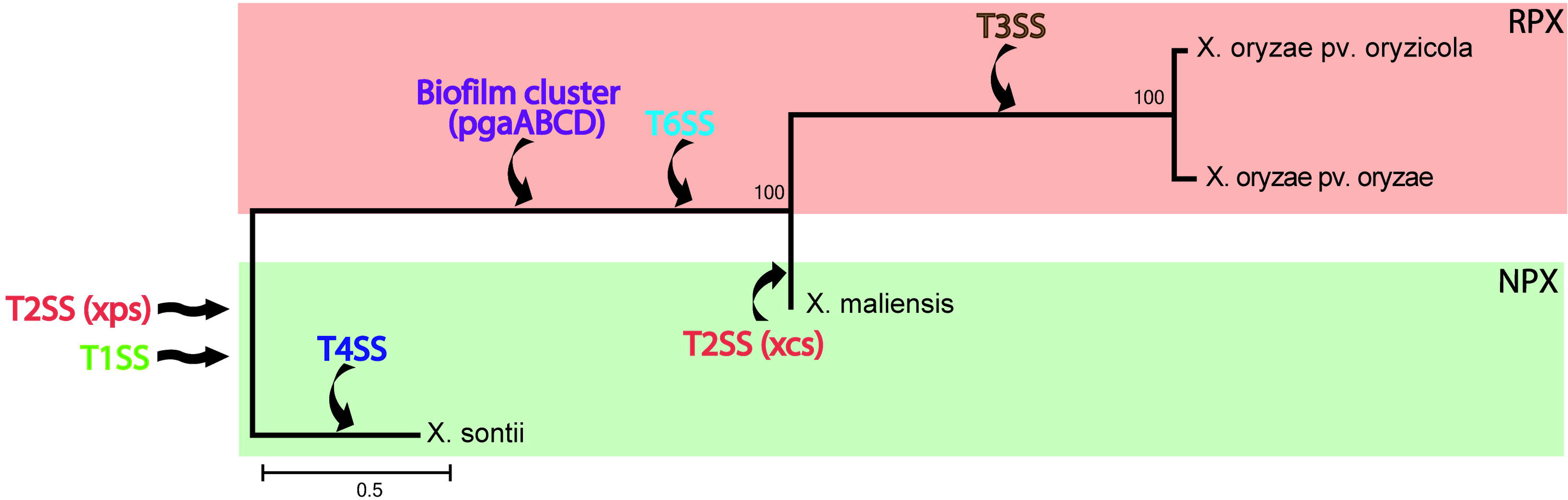
**Pattern of acquisition or loss of pathogenicity** related clusters among *X. oryzae* (in red box) and *X. sontii* and *X. maliensis* (in green box). Black arrows depict the gene flow.

On the other hand, among the two T2SS (*xps* and *xcs*), *X. sontii* and all the *X. oryzae* isolates were having *xps*. Presence of an additional non-canonical T2SS *xps* cluster in *X. maliensis* but absence in all other strains indicate on-going dynamics and selection within the species. Further, cell wall degrading enzymes (secreted by T2SS) are also present in all the strains under study, though with differential repertoires. Conservation of T2SS and most of the effectors where we further see redundancy suggests that they were present in ancestor of both the lineages and were acquired for plant adaptation. The exceptions are *cbsA* (exclusively present in *X. oryzae* except for *X. oryzae* pv. oryzicola) and *cel9A* (exclusively present in *X. sontii* and *X. maliensis*) genes encoding cellobiosidase and cellulose respectively suggesting their specific acquisition. In an earlier study, cellobiosidase gene is reported to be a major virulence factor (Jha, Rajeshwari et al. 2007).

Apart from protein secretion systems, adhesion to the host is the most crucial step for bacteria irrespective of the lifestyle followed. However, redundancy is reported in this case as well and mutations in majority of these genes do not completely abolish virulence (Das, Rangaraj et al. 2009). However, mutation in *yapH* encoding gene is reported to abolish virulence of *Xanthomonas* due to deficiency in leaf attachment and entry (Das, Rangaraj et al. 2009). In the present study, exclusive presence of *yapH* in *X. oryzae* suggests its importance in emergence of pathogenic strains of *Xanthomonas* in rice. Conservation of EPS biosynthetic pathway, siderophore biosynthetic gene clusters, iron uptake genes and regulatory genes (*xibR*) and other well-known genes like *rpf* cluster, several two component systems for mediating environmental signals and *gigX* cluster for flagellar glycosylation (Yu, Chen et al. 2018) among all the rice associated *Xanthomonas* point to their primary role in plant adaptation per se. At the same time, this study also allowed us to track genes that are implicated in virulence but are under reductive evolution in NPX. For example, the T1SS related *raxX* cluster that encodes a peptide recognized by Xa21 rice resistance gene is present in all pathogenic strains but seem to be under reductive evolution in non-pathogenic strains. It indicates that strategies once suitable for adaptation which have been hijacked or exploited by pathogenic counterparts but are being lost from the non-pathogenic counterparts. Hence, such plant adaptation genes which are now evolved to be virulence genes can be important candidates for genetic studies of pathogenic isolates of rice.

It is interesting to note that *X. maliensis* is a non-pathogenic strain lacking T3SS and its effectors, but on the other hand, it contains a T6SS, biofilm formation cluster (*pgaABCD)* and *raxX, raxST, rasA* genes of T1SS and does not have T4SS all of which are features associated with the pathogenic strains. Furthermore, it is found to be phylogenomically more related to pathogenic than to non-pathogenic isolates. This might be representing a second wave of evolution of non-pathogenic isolates, which remains non-pathogenic but have some of the genes associated with the pathogenic lineage. What do these intermediate strains represent? Do they represent variations of a non-pathogenic lifestyle that benefit from acquisition of genes that are normally present in pathogenic strains? Alternatively, could they represent non-pathogenic strains that have set off on the path towards pathogenicity? Further studies on these kinds of intermediate strains as well as the non-pathogenic strains can lead to a better understanding of the mechanisms of adaptation of the genus *Xanthomonas* on plants.

### Conclusion

Although the genus *Xanthomonas* has evolved as highly successful plant pathogens, this study highlights the scope and possibilities for detailed comparatives studies on both pathogenic and non-pathogenic *Xanthomonas* strains that are associated with a particular plant. A small core genome shared amongst all rice associated *Xanthomonas* strains and non-uniformity of various horizontally acquired gene clusters suggest a non-recent divergence of these lineages and occurrence of various events of gene gain and gene loss amongst these rice associated bacteria. Interestingly the study has revealed many well characterized gene clusters that are common to both life-styles which are indicative of a role for these functions in plant adaptation. Moreover, the study has revealed many novel and interesting gene(s) that are unique to each of the groups. These can be promising material for researchers engaged in genetic and molecular studies to understand *Xanthomonas* biology in particular and pathogenesis in general.

## Methods

### Genome sequencing, assembly and annotation

Genomic DNA extraction was carried out using ZR Fungal/Bacterial DNA MiniPrep kit (Zymo Research, Irvine, CA, USA). Qualitative assessment of DNA was performed using NanoDrop 1000 (Thermo Fisher Scientific, Wilmington, DE, USA) and agarose gel electrophoresis. Quantitative test was performed using Qubit 2.0 fluorometer (Life Technologies). Nextera XT sample preparation kits (Illumina, Inc., San Diego, CA, USA) were used to prepare Illumina paired-end sequencing libraries (250 × 2 read length) with dual indexing adapters. In-house sequencing of the Illumina libraries was carried out on Illumina MiSeq platform (Illumina, Inc., San Diego, CA, USA). Adapter trimming was performed automatically by MiSeq control software (MCS), and remaining adapters were detected by NCBI server and were removed by manual trimming. Sequencing reads were *de novo* assembled into high quality draft genome on CLC Genomics Workbench v7.5 (CLC bio, Aarhus, Denmark) using default settings. Genome annotation was performed by NCBI PGAP pipeline (http://www.ncbi.nlm.nih.gov/genome/annotation_prok).

### Phylogenomic and taxonogenomic analysis

Phylogenomic analysis based on more than 400 putative conserved genes was constructed using PhyloPhlAn (Segata, Börnigen et al. 2013) from whole genome proteome data. Ortholog searching, multiple sequence alignment and phylogenetic construction were done using USEARCH v5.2.32 (Edgar 2010), MUSCLE v3.8.31 (Edgar 2004) and FastTree v2.1 (Price, Dehal et al. 2009) respectively. All strains under study were used as an input to PhyloPhlAn. *Stenotrophomonas maltophilia* was used as an outgroup. Taxonogenomic analysis of all the strains carried out using orthoANI values calculated by using USEARCH (Edgar 2010) for taxonomic status of bacterial species.

### Virulence related gene clusters

Various genes clusters related to virulence (supplementary table 1) examined in the present study were retrieved from NCBI. The tBLASTn searches were performed using retrieved sequences as query. Cut-off for similarity was set to be 40% and coverage was 50%. Cluster figures were generated using Easyfig v2.2.2(Sullivan, Petty et al. 2011) and heat maps for the blast results were generated using GENE-Ev3.0.215 (https://www.broadinstitute.org/).

### Lineage-wise gene content analysis

Lineage-wise core genes were fetched from gene profiling done by roary pan genome pipeline (Page, Cummins et al. 2015). Since, strains we were dealing with were from three different species (*X. oryzae, X. maliensis* and *Xanthomonas* sp.), pan genome analysis was performed using identity cut-off of 60%. Further, GC content of genes were calculated and genes with atypical GC content were identified (GC content not in range of 64.5 +-2%). Functional analysis of the lineage specific core genes was performed using EggNOG, Prokka and GenBank annotation was also considered.

### Biofilm assay

Cultures were grown in NB (nutrient broth) media at 28 °C at 180 rpm until OD at 600 nm reaches 0.5. Cultures were then diluted 1:10 with NB media containing 2% glucose. Then, 1ml of diluted culture was dispensed into borosilicate glass tubes. Test tubes were then incubated at 28 °C for 7 days. After 7 days, media was discarded and non-adherent cells were removed by washing three times with sterile water. Quantification of biofilm formation was assessed by crystal violet (CV staining. Briefly, biofilms were fixed at 60 °C for 20 min, and stained with 1.5ml of 0.1% CV for 45 min. The dye was discarded, and the plate was rinsed in standing sterile water and then allowed to dry for 30 min at RT. Stained biofilms were dissolved in 1.5ml of ethanol: acetone (80:20) and finally OD was read at 595nm using spectrophotometer.

### Recombination and mutation analysis

Impact of recombination and mutations was evaluated among the strains under study using ClonalFrameML analysis. Core genome alignment was performed by MAUVE v2.3.1(Darling, Mau et al. 2004) and it was further used to generate maximum likelihood tree by PhyML v3.1(Guindon, Dufayard et al. 2010).These alignment and tree were further subjected to ClonalFrameML analysis (Didelot and Wilson 2015) to calculate frequency and impact of recombination and mutation.

## Author Contributions

SM and KB performed strain isolation, identification and genome sequencing. KB performed phylogenomic and comparative analysis. KB and SK have performed virulence gene cluster analysis. AK performed biofilm assay. KB have drafted manuscript with inputs from SK, SM, AK, RVS and PBP. PBP conceived the study and participated in its design with KB, SM and RVS. All the authors have read the manuscripts and approved the manuscript.

## Acknowledgements

KB is supported by fellowship of University Grant Commission (UGC). SM is supported by fellowship of Council of Scientific and Industrial Research (CSIR). AK is supported by DST-INSPIRE fellowship. This work is supported by a project entitled “Plant-Microbe and Soil Interactions” BSC0117 of Council of Scientific and Industrial Research (CSIR) to PBP.

---

**Supplementary table 1:** Information regarding virulence clusters used for the analysis.

